# High interictal connectivity within the resection zone is associated with favorable post-surgical outcomes in focal epilepsy patients

**DOI:** 10.1101/459008

**Authors:** Preya Shah, John Bernabei, Lohith Kini, Arian Ashourvan, Jacqueline Boccanfuso, Ryan Archer, Kelly Oechsel, Timothy H. Lucas, Danielle S. Bassett, Kathryn A. Davis, Brian Litt

## Abstract

**Objective:** Patients with drug-resistant focal epilepsy are often candidates for invasive surgical therapies. In these patients, it is necessary to accurately localize seizure-generators to ensure seizure freedom following intervention. While intracranial electroencephalography (iEEG) is the gold standard for mapping networks for surgery, this approach requires inducing and recording seizures, which may cause patient morbidity. The goal of this study is to evaluate the utility of mapping interictal (non-seizure) iEEG networks to identify targets for surgical treatment.

**Methods:** We analyze interictal iEEG recordings and neuroimaging from 27 focal epilepsy patients treated via surgical resection. We generate interictal functional networks by calculating pairwise correlation of iEEG signals across different frequency bands. We identify electrodes falling within surgically resected tissue (i.e. the resection zone), and compute node-level and edge-level synchrony in relation to the resection zone. We associate these metrics with post-surgical outcomes.

**Results:** Greater overlap between resected electrodes and highly synchronous electrodes is associated with favorable post-surgical outcomes. Additionally, good outcome patients have significantly higher connectivity localized within the resection zone compared to those with poorer postoperative seizure control. This finding persists following normalization by a spatially-constrained null model.

**Conclusions:** This study suggests that spatially-informed interictal network synchrony measures can distinguish between good and poor post-surgical outcomes. By capturing clinically relevant information during interictal periods, our method may ultimately reduce the need for prolonged invasive implants and provide insights into the pathophysiology of an epileptic brain. We discuss next steps for translating these findings into a prospectively useful clinical tool.

## 1. Introduction

Epilepsy is a common neurological disorder that affects over 50 million people worldwide, ^1^ one-third of whom experience uncontrolled seizures despite medication.^2^ Within this group, approximately 80% have localization-related seizures and are candidates for surgical removal of the seizure-generating region in the brain.^3^ Accurate seizure localization is needed in these patients to maximize chances of seizure freedom and minimize deficits following surgery. With the recent development of more targeted alternatives to surgery, such as laser ablation^4^ and neurostimulation through implantable devices^5^, precise localization is becoming increasingly necessary to guide therapy.

Intracranial EEG (iEEG) is currently the gold standard for localizing seizures, particularly in patients without clear lesions on clinical imaging.^6^ In this approach, implanted subdural and depth electrodes record brain signals for up to several weeks, with the intent of capturing ictal events and identifying seizure onset regions. While iEEG can record seizures at high spatial and temporal resolution, it has important limitations. For example, seizures are often provoked during the recording period via medication withdrawal and sleep deprivation; in some patients, these provoked seizures may be fundamentally different from stereotypical spontaneous seizures, with the potential to misinform localization attempts.^7^ Additionally, seizure provocation and prolonged implantation while waiting to record seizures increases the risk of complications such as infection, deep vein thrombosis, musculoskeletal injury, and postictal psychosis.^8,9^ In some cases, seizures may not occur during the implant period, rendering the study inconclusive. Prolonged hospitalizations with multiple surgeries – usually at least one for electrode implantation and one for removal – add greatly to patient inconvenience and consumption of limited resources. Eliminating the need to record seizures would be a marked advance in improving presurgical evaluation for epilepsy.^10,11^ In the long term, a procedure in which interictal recording, stimulation mapping, and subsequent intervention could take place in a single session, comparable to cardiac electrophysiology, would have notable benefits compared to current procedures.

Recent evidence indicates that seizures most commonly arise from abnormal brain networks rather than isolated focal lesions.^12,13^ Therefore, in order to accurately map seizure generation, it is important to identify brain network abnormalities in epilepsy. Functional networks derived from correlations between iEEG signals show promise in highlighting seizure onset networks, distinguishing between focal and generalized seizures, and predicting outcomes.^14–16^ While most previous iEEG network studies analyze ictal and preictal data, recent evidence suggests that interictal recordings are also informative for localizing epileptic networks.^17^ This notion is further supported by recent studies demonstrating that ictal and interictal iEEG network subgraphs are topologically similar, and that patterns of high frequency activity propagation during seizures are recapitulated interictally.^18,19^ Moreover, epileptic brain networks are fundamentally altered, as reflected by cognitive deficits and imaging abnormalities, in many patients in regions associated with seizures.^20–23^ These findings suggest that iEEG can provide valuable information without the need to capture seizure events.

In this study, we evaluate the utility of interictal network analysis in mapping seizure networks. While the ground-truth identity of seizure-generating networks is inherently unknown, rigorously quantified information about the surgical resection zone combined with outcome data can serve as a valuable proxy. Namely, if a patient has a good post-surgical outcome, a reasonable assumption is that vital parts of the seizure network are contained in the resection zone. In contrast, in poor outcome patients, the resection zone likely did not include critical regions of the seizure-generating network. Therefore, we characterize network connectivity inside and outside of the resection zone in good and poor outcome patients. We hypothesize that patients with highly synchronous nodes removed are more likely to have good outcomes. We also propose that this work can further our understanding of functional network topology in the epileptic human brain, and ultimately reduce the need for prolonged implant times and resulting patient morbidity.

## 2. Methods

### 2.1 Subjects

We retrospectively studied 27 adult patients undergoing pre-surgical evaluation for drug-resistant epilepsy at the Hospital of the University of Pennsylvania and at the Mayo Clinic. All patients presented with focal onset seizures and were subsequently treated by surgical resection, with at least 1-year post-surgical outcomes as measured by Engel classification score and/or ILAE criteria. Patients were divided into two groups: good outcome (Engel I or ILAE 1-2) and poor outcome (Engel II-IV or ILAE 3-6). All patients gave consent to have their anonymized iEEG data publicly available on the International Epilepsy Electrophysiology Portal (www.ieeg.org).^24,25^ Clinical and demographic information is available in **Table 1**.

**Table 1:** Clinical and demographic patient information. Legend – L: Left; R: Right; TL: Temporal Lobe; FL: frontal lobe, FPL: fronto-parietal lobe, FTL: fronto-temporal lobe, MTS: mesial temporal sclerosis; MCD: malformation of cortical development; N/A: not available.

| Patient # | Age | Sex | Resected Region | Outcome |
| --- | --- | --- | --- | --- |
| 1 | 21 | M | LFL | Engel 1D |
| 2 | 37 | M | RTL | Engel 1B |
| 3 | 28 | F | RTL | Engel 1A |
| 4 | 33 | M | LFPL | Engel 1B |
| 5 | 40 | M | RFL | Engel 1C |
| 6 | 25 | F | LTL | Engel 1A |
| 7 | 57 | F | LTL | Engel 2D |
| 8 | 54 | M | LTL | Engel 2A |
| 9 | 41 | F | LTL | Engel 2B |
| 10 | 56 | F | RTL | Engel 1A |
| 11 | 29 | M | LPL | Engel 2A |
| 12 | 25 | F | LTL | Engel 1C |
| 13 | 24 | M | LFL | Engel 1D |
| 14 | 35 | F | LTL | Engel 1D |
| 15 | 48 | F | RTL | Engel 1B |
| 16 | 39 | M | RTL | Engel 1A |
| 17 | 45 | F | LTL | Engel 1B |
| 18 | 36 | M | RTL | Engel 1A |
| 19 | 40 | F | RTL | Engel 1B |
| 20 | N/A | M | RFL | ILAE1 |
| 21 | N/A | F | RFTL | ILAE4 |
| 22 | N/A | M | RTL | ILAE4 |
| 23 | N/A | M | LTL | ILAE5 |
| 24 | N/A | M | RFL | ILAE5 |
| 25 | N/A | F | LTL | ILAE5 |
| 26 | N/A | M | LFPL | ILAE4 |
| 27 | N/A | F | RTL | ILAE5 |

### 2.2 Intracranial EEG acquisition

Cortical surface and depth electrodes were implanted in patients based on clinical necessity. Electrode configurations (Ad Tech Medical Instruments, Racine, WI) consisted of linear cortical strips and two-dimensional cortical grid arrays (2.3 mm diameter with 10 mm inter-contact spacing), and linear depths (1.1 mm diameter with 10 mm inter-contact spacing).

Continuous iEEG signals were obtained for the duration of each patient’s stay in the epilepsy monitoring unit. For each subject, we obtained one clip of interictal data consisting of the first 6 hours of artifact-free recording at least 4 hours removed from any seizure event. Seizure events were labeled by a board-certified epileptologist and were checked for consistency with clinical documentation.

### 2.3 Electrode and resection zone localization

All patients underwent a clinical epilepsy neuroimaging protocol. Pre-implant T1-weighted MPRAGE MRI and post-implant CT images were acquired in order to localize electrodes within or on the surface of each patient’s brain. Furthermore, patients underwent a post-resection imaging protocol acquired between 6-8 months after resection, which consisted of T1-weighted MPRAGE MRI and axial FLAIR MRI sequences. Images were anonymized and coregistered to each patient’s pre-implant T1 MRI space for localization and segmentation.

Electrodes were identified via thresholding of the CT image and labeled using a semiautomated process. For all subjects, all images were registered to the pre-implant MRI space using 3D rigid affine registration, with mutual information as the similarity metric. Pre-implant MR images were diffeomorphically coregistered to post-resection MR images to quantitatively identify the resection zone. Resection zones were segmented semi-automatically via the ITK-SNAP random forest classifier feature using coregistered MPRAGE and FLAIR imaging. The resection zone was dilated by 5% of the iEEG network in order to mimic effects of gliosis and scarring adjacent to surgically removed tissue.^26^ Using these resected regions along with the electrode localizations, we determined the identities of the electrodes present in the resection zone. Coregistration steps utilized Advanced Normalization Tools (ANTs) software.^27,28^

### 2.4 Functional network analysis

Following removal of electrode channels obscured by artifact (marked by a board-certified epileptologist), interictal iEEG clips were common-average referenced to reduce potential sources of correlated noise.^29^ Each clip was then divided into 1 s non-overlapping time windows in accordance with previous studies.^14,19,30,31^ To generate a network representing broadband functional interactions between iEEG signals for each 1 s time window, we employed a method described in detail previously.^19^ Signals were notch-filtered at 60 Hz to remove power line noise, low-pass and high-pass filtered at 115 Hz and 5 Hz to account for noise and drift, and pre-whitened using a first-order autoregressive model to account for slow dynamics. Functional networks were then generated by applying a normalized cross-correlation function between the signals of each pair of electrodes within each time window. Next, to gain an understanding of iEEG networks across different frequencies, we generated functional networks across physiologically relevant frequency bands as described in detail in a previous study.^14^ Specifically, multitaper coherence estimation was used to compute functional coherence networks for each 1 s window across four frequency bands: alpha/theta (5-15 Hz), beta (15-25 Hz), low-gamma (30-40 Hz), and high-gamma (95-105 Hz). Both broadband and frequency-specific networks were represented as full-weighted adjacency matrices. In this model, each electrode serves as a node of the network, and measurements of connectivity between pairs of electrodes serve as edges. The networks were averaged across the full 6 hours to obtain one functional network for each patient for each frequency band. A schematic of this pipeline is provided in **Figure 1**.

**Figure 1:**
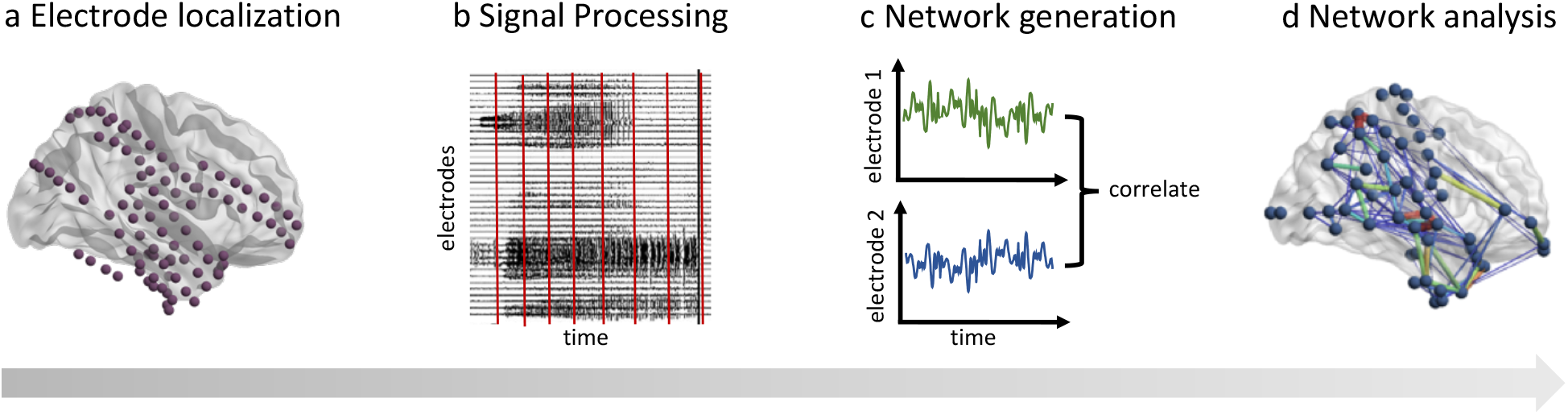
Schematic of subject-level iEEG network analysis pipeline. (a) Using structural imaging. the location of each electrode is identified on the brain surface and within the parenchyma. (b) Interictal iEEG signals are processed and divided into 1 s windows. (c) For each 1 s window, a broadband functional connectivity network is generated by calculating the correlation between iEEG signals across electrode pairs. Frequency-specific networks are similarly constructed by calculating coherence between iEEG signals measured by electrode pairs. (d) Node-level and edge-level network analyses are computed on these resulting networks, in relation to the resection zone.

To quantify the degree of synchrony of each node in the network, we computed the nodal strength, defined for each node as the sum of the weights of all edges connected to that node.^32^–^33^ We defined “highly synchronous nodes” to be nodes with a value of strength that is at least 1 *z-* score above the mean. Next, we defined the *strength selectivity* of the resection zone as the spatial overlap between the nodes within the resection zone and the highly synchronous nodes. Overlap was computed using the Dice Similarity Coefficient (DSC), which ranges from 0 to 1 and is defined as 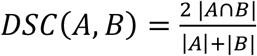 where *A* and *B* are two binary sets. We compared the strength selectivity for good and poor outcome patients across all frequency bands, and we repeated the analysis for *z* thresholds ranging from 0 to 2. Furthermore, to assess what types of connections were contributing to the observed differences in strength selectivity in good vs. poor outcome patients, we delineated the following three edge types: (i) connections between nodes within the resection zone (RZ-RZ), (ii) connections between one node within the resection zone and one node outside the resection zone (RZ-OUT), and (iii) connections between nodes outside the resection zone (OUT-OUT). For each subject, we computed the mean edge weights within each of these categories. We compared the mean edge weights among these three categories within both good and poor outcome patients. Furthermore, we computed differences in these categories between the two patient groups.

Given that neighboring electrodes are more likely to be highly correlated due to spatial proximity or due to common source measurements, we generated a spatially-constrained null model. For each patient with *N* resected electrodes, we sampled clusters of *N* spatially contiguous electrodes, using Euclidean distance to determine the closest electrodes. We repeated the edge-weight analysis after normalizing the RZ-RZ, RZ-OUT, and OUT-OUT edge weights by the null distribution of edge weights for each category. Normalization was carried out by subtracting the mean and dividing by the standard deviation of the null values. Non-parametric Mann-Whitney *U* tests were used for all pairwise comparisons. Our code is available at http://github.com/shahpreya/Epimapper.

## 3. Results

We constructed spatial maps of nodal strength, along with the overlaid resection zone, for each individual patient (**Figure 2**). At the group level, we observed significantly higher broadband and beta strength selectivity in good outcome patients vs. poor outcome patients, using a *z* threshold of 1 (*p* < 0.05, Mann-Whitney *U* test) (**Figure 3A**). Sweeping across a range of z thresholds from 0 to 2 revealed a trend of higher strength selectivity in good vs. poor outcome patients in all frequency bands, with significant differences for broadband (*z* = 1), beta band (*z* = 0.5 to *z* = 1.25) and low-gamma band (*z* = 1.75 to *z* = 2) (**Figure 3B**).

**Figure 2:**
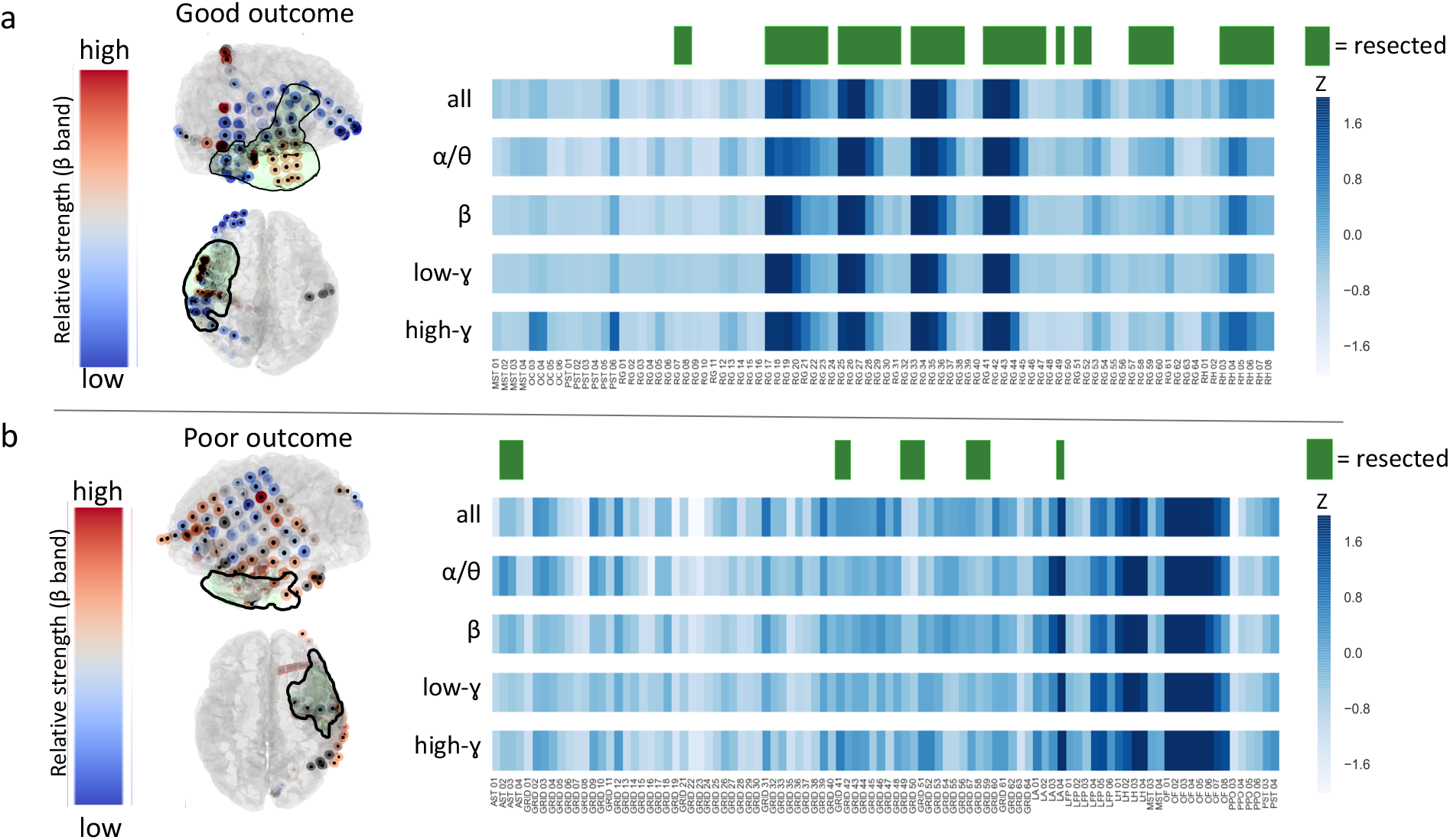
Patient-level strength selectivity analysis. For an example good outcome patient (a) and poor outcome patient (b), we provide spatial maps of nodal strength in the beta band only, along with corresponding 2D heat maps of nodal strength in all frequency bands. Resection zones are highlighted in green.

**Figure 3:**
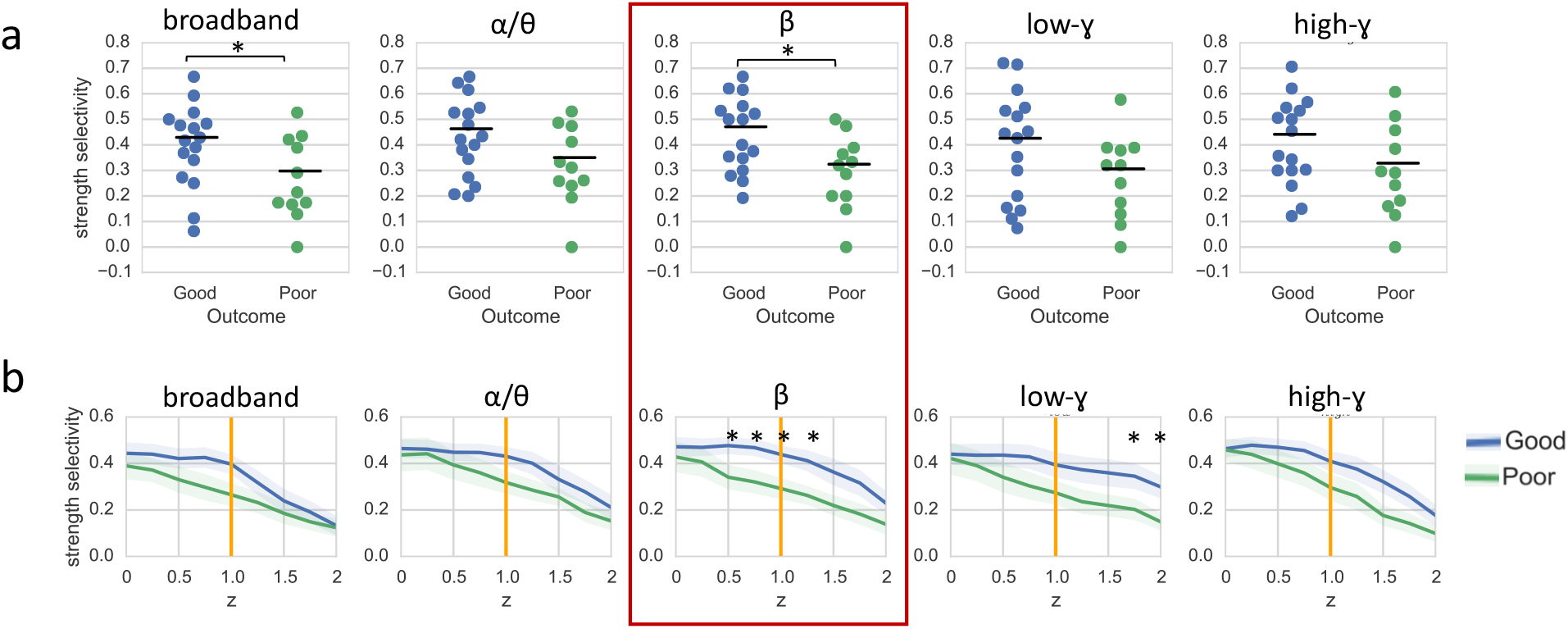
Group-level strength selectivity analysis, (a) Strength selectivity in all tested frequency bands with a *z* threshold of 1 reveals significantly higher broadband and beta strength selectivity in good outcome patients vs. poor outcome patients. (b) A sweep across multiple *z* thresholds from 0 to 2 reveals significant outcome-dependent differences in strength selectivity for broadband (*z* = 1), beta band (*z* = 0.5 to *z* = 1.25) and low-gamma band (*z* = 1.75 to *z* = 2) networks (mean +/-standard error). Beta band strength selectivity distinguishes between good and poor outcome patients across the widest range of *z* thresholds (red box). *\*p* < 0.05, Mann-Whitney *U* test.

Next, we sought to understand whether observed strength selectivity findings were due to connectivity within the resection zone, or connectivity between the resection zone and extra-resection regions. Since strength-selectivity findings were most prominent in the beta band, we focused our edge-level analysis on the beta band networks; to assess sensitivity and specificity, we also repeated the analysis across all frequency bands. We found that connections within the resection zone (RZ-RZ) were significantly stronger than RZ-OUT and OUT-OUT connections. We also observed that RZ-RZ connections were stronger in good outcome patients than in poor outcome patients (*p* < 0.05) (**Figure 4**). Notably, these findings persisted across all frequency bands (*p* < 0.05). After normalization by a spatially constrained null model, RZ-RZ connections were still significantly stronger in good outcome patients than in poor outcome patients (*p* < 0.05). While this trend was present in all frequency bands, it was statistically significant for beta, low-gamma, and high-gamma bands. Additionally, in both good and poor outcome patients, normalized RZ-RZ connections were stronger than RZ-OUT connections, and RZ-OUT connections were stronger than OUT-OUT connections, with the additional finding of RZ-RZ > RZ-OUT in good outcome patients (*p* < 0.05). Notably, these findings also persisted across all frequency bands.

**Figure 4:**
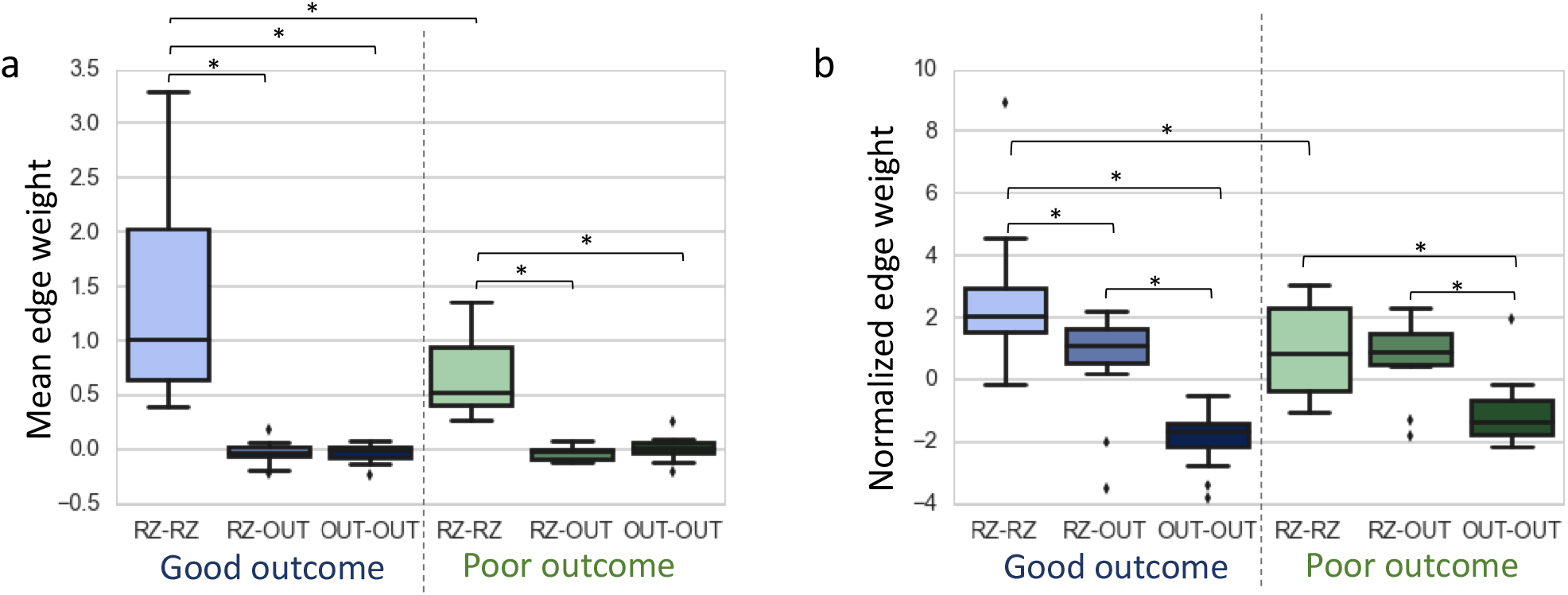
Edge-level analysis in relation to resection zone, shown for the beta frequency band. (A) Connections within the resection zone (RZ-RZ) are significantly stronger than RZ-OUT and OUT-OUT connections, and RZ-RZ connections are stronger in good outcome patients than in poor outcome patients. (B) After normalization by a spatially-constrained null model, RZ-RZ connections remain significantly stronger in good outcome patients than poor outcome patients. Additionally, in both good and poor outcome patients, normalized RZ-RZ connections are stronger than RZ-OUT connections and RZ-OUT connections are stronger than OUT-OUT connections. Finally, we also observe that RZ-RZ connections are stronger than RZ-OUT connections in good outcome patients in comparison to poor outcome patients. *\*p* < 0.05, Mann-Whitney *U* test.

Given similar findings across different frequency bands, we sought to directly probe the similarity of the frequency-specific function networks. Therefore, we calculated the correlation coefficient between the edges of the mean functional networks for each pair of frequency bands, across all subjects. We found a high degree of correlation across these networks (r = 0.95-0.91), with the highest correlations between neighboring frequency bands (e.g., alpha-theta vs. beta: r = 0. 91), and the lowest correlations being observed between bands with larger frequency separation (e.g., alpha-theta vs. beta: r = 0.75) **(Figure 5**).

**Figure 5:**
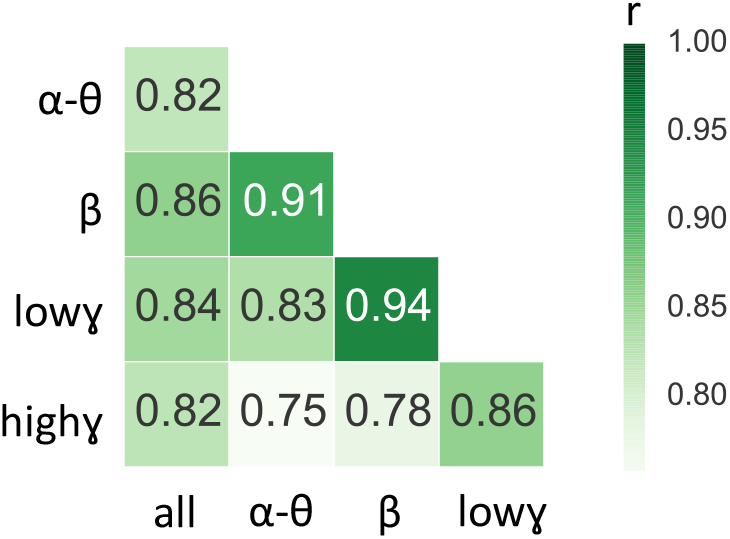
Matrix of similarity values between functional networks generated using different frequency bands. Similarity values were obtained by measuring the Pearson correlation coefficient between edges in each pair of networks for each subject, and then by averaging these correlation coefficients across subjects. We observe strong correlations across all pairs of networks (r = 0.75-0.91), with the highest correlation coefficients being observed between neighboring frequency bands and the lowest correlation coefficients being observed between bands with larger frequency separation.

## 4. Discussion

In this study, we evaluate the association of interictal network synchrony within the resection zone with post-surgical outcomes in drug-resistant focal epilepsy patients. We determine that high interictal strength selectivity is associated with better outcomes. This effect appears to be driven largely by connectivity within the resection zone. Our findings suggest that interictal recordings can provide valuable information to identify putative seizure-generating networks. Employing quantitative tools on early interictal recordings can maximize information gained from iEEG recordings while significantly reducing recording times.

### 4.1 High interictal connectivity within the resection zone is associated with good outcomes

We define the strength selectivity of the resection zone as a simple measure of overlap between electrodes within the resected region and highly synchronous electrodes. We find that strength selectivity is higher in good outcome patients compared to poor outcome patients. The notion that removing highly synchronous nodes would lead to favorable outcomes is consistent with our understanding that epilepsy is characterized by abnormal hypersynchronous neuronal firing.^34^ We demonstrate the utility of our method using the simple, non-parametric measure of node strength, and we explore our findings across a range of physiologically relevant frequency bands.

Although we observe a trend of increased strength selectivity in good outcome patients across a range of *z* thresholds using both broadband and frequency-specific networks, the finding is most significant in functional networks constructed in the beta frequency band. While beta frequency oscillations are thought to be associated with long-range communication between regions,^35^ the role of oscillatory activity across different frequency bands is still complex, with known interactions between different frequencies.^36–38^ Moreover, our direct analysis of network similarity across frequency bands indicates that the frequency-specific networks are highly correlated with each other. This high similarity may be due to our data processing pipeline, as we compute average networks across six-hour periods, a choice which was motivated by our interest in extracting information from stable functional networks. Kramer *et al.* have shown that while functional iEEG networks are highly variable when estimated from time windows spanning a few seconds, stable network topology emerges when using time windows of 100 seconds or more and persists across frequency bands.^39^ Our group has previously shown that long-term interictal functional network connectivity across all frequency bands can accurately predict structural connectivity derived from white-matter tractography. Therefore, it is possible that long-term interictal functional networks are highly similar across different frequency bands because these functional networks echo the underlying structural connectome that gives rise to wideband functional dynamics.^40^

Our findings contribute to a growing body of recent work aiming to identify epileptogenic networks and predict outcomes based on node-level measures.^15,17,41–45^ While most of this prior work uses ictal or preictal recordings, we focus on deriving information from the earliest available interictal data. Moreover, unlike the majority of previous studies, we apply quantitative imaging methods and semi-automated segmentation techniques to delineate the resection cavity, rather than utilizing subjective clinical identification of electrodes that are resected or part of the seizure onset zone. Marking the seizure onset zone is an inexact process; moreover, seizure onset zone areas may be omitted from the resection zone due to practical concerns (e.g. proximity to blood vessels or eloquent cortex). Therefore, it may be advantageous, and more direct, to determine outcomes based on what was actually removed rather than what was identified to be part of the seizure onset zone.

Our edge-level analysis reveals that the majority of network synchrony is attributable to intra-resection connections. This finding is similar to previous analyses illustrating that connectivity within the seizure onset zone is higher than both connectivity outside of the seizure onset zone and connectivity bridging seizure onset and non-seizure onset regions.^31,46,47^ We find higher intra-resection zone connectivity in good vs. poor outcome patients. These analyses suggest that seizure-generating regions are functionally distinct or isolated in some way from surrounding brain regions in focal epilepsy patients, and that removal of these functionally isolated regions improves the likelihood of a successful outcomes.

Our finding that good-outcome patients had higher intra-resection zone connectivity than poor-outcome patients persists following normalization by a spatially-constrained null model. Previous similar studies have generated null distributions by sampling *N* random electrodes from the network, where *N* is the number of electrodes in the region of interest (e.g. the resection zone).^17,41^ However, these models do not consider two key facts: (1) that neighboring electrodes are more likely to have higher functional connectivity due to structural connections or common source signals, and (2) that surgical practice necessitates removal of spatially contiguous brain regions rather than distant, randomly distributed electrodes. Our spatially constrained null model therefore provides a more realistic set of random resections with which to normalize our connectivity findings. Since this spatially constrained null is more stringent than a random resection-based null, it is likely that our intra-resection connectivity findings are biologically significant and not simply due to spatial proximity.

### 4.2 Methodological Considerations and Limitations

One concern inherent to all iEEG data analysis is that the entire brain is not sampled, as electrode locations are based on clinical necessity. While spatial coverage is sparse to minimize patient morbidity, electrodes are placed with the intent of capturing regions hypothesized to be part of the seizure network. Therefore, the seizure network should still be captured, particularly in patients with good outcomes. However, it is possible that the seizure network is not adequately covered, particularly in poor-outcome patients. Moreover, the spatial distribution and number of nodes in the network may impact the topological properties derived from the network. Recent efforts to map whole-brain iEEG may help circumvent this issue.^40,48,49^ Furthermore, source modeling from high-density scalp EEG or MEG recordings, as well as data from functional and structural neuroimaging such as MRI and PET, could complement our intracranial analysis and allow spatial sampling of the whole brain. This work remains in progress and will require rigorous quantitative validation.

Another limitation of this study is that the resection cavity may include more tissue than was necessary to resect. For example, patients with temporal lobe epilepsy often have a standard anterior temporal lobectomy that removes both temporal neocortex and mesial temporal structures, even if only one of those areas is involved in the epileptic network.^50^ This fact does not invalidate our results; rather it provides an opportunity to refine them. Recent increases in focal laser ablation and neurostimulation approaches in the United States will allow us to test the method on these more targeted interventions and compare to prior standard approaches involving larger volume tissue resections. These studies are currently underway.

We demonstrate a framework for mapping interictal functional networks using simple measures of network synchrony on a moderately sized dataset. In order to bring this framework to clinical practice, the next step is to generate a suite of multimodal network-based features and assess the capacity of these features to predict candidate targets for surgical removal, using large multi-institutional datasets. By sharing our data, code, and analysis approach, we hope to facilitate translation of quantitative seizure-mapping tools to clinical practice.

### 4.3 Future Directions

It is important to consider what will be required to translate studies such as ours from retrospective computational experiments to clinical utility. As noted above, the first step is validation on a much larger group of patients, ideally representative of the cross-section of patients who undergo epilepsy surgery and including those treated more focally and those with larger resections. The recent significant advances in our understanding of both ictal and interictal network dynamics and their relationship to epilepsy surgical clinical outcomes at a group level suggests that these investigations will lead to patient specific outcome prediction. Machine learning approaches carried out on a large, multimodal dataset, including structural and functional neuroimaging and iEEG recordings, holds promise for identifying the features predictive of individual patient outcomes and for translating our work to be tested in a prospective clinical trial. The ultimate goal, of course, is that this body of work results in either: (1) network-assisted procedures, in which standard procedures might be augmented with ablation or resection of additional electrodes not originally intended to be removed in more standard procedures, or (2) complete guidance of resection or ablation by network models. If effective, the hope is that epilepsy surgical interventions might eventually be done, based upon successful trials of interictal network modeling guidance for surgery, in a single session, both recording and ablation or resection, rather than requiring two weeks of hospitalization and multiple surgical procedures.

### 4.4 Conclusion

We demonstrate that high interictal connectivity within the resection zone is associated with favorable post-surgical outcomes in drug-resistant focal epilepsy patients. This study is one of a series of investigations that we hope will lead to translation of our methods into clinical practice, ultimately allowing for automated and optimized seizure localization and treatment with reduced need for prolonged invasive implants.

## 5. Acknowledgements

This work was supported by National Institutes of Health grants 1R01NS099348 and K23-NS073801. We also acknowledge support by the Thornton Foundation, the Mirowski Family Foundation, the ISI Foundation, the John D. and Catherine T. MacArthur Foundation, the Sloan Foundation, the Paul Allen Foundation, Neil and Barbara Smit, and Jonathan Rothberg.

